# Data science identifies novel drug interactions that prolong the QT interval

**DOI:** 10.1101/024745

**Authors:** Tal Lorberbaum, Kevin J. Sampson, Raymond L. Woosley, Robert S. Kass, Nicholas P. Tatonetti

## Abstract

Drug-induced prolongation of the QT interval on the electrocardiogram (long QT syndrome, LQTS) can lead to a potentially fatal ventricular arrhythmia called *torsades de pointes* (TdP). 180 drugs with both cardiac and non-cardiac indications have been found to increase risk for TdP, but drug-drug interactions contributing to LQTS (QT-DDIs) remain poorly characterized. Traditional methods for mining observational healthcare data are poorly equipped to detect QT- DDI signals due to low reporting numbers and a lack of direct evidence for LQTS. In this study we present an integrative data science pipeline that addresses these limitations by identifying latent signals for QT-DDIs in the FDA’s Adverse Event Reporting System and retrospectively validating these predictions using electrocardiogram data in electronic health records. We present 26 novel QT-DDIs flagged using this method that warrant further investigation.

**Key Points:**

- Drug-induced long QT syndrome (LQTS) can lead to potentially fatal arrhythmias (*torsades de pointes*, TdP). Drug-drug interactions that prolong the QT interval (QT- DDIs) can be clinically significant but remain poorly characterized.
- Observational health data (such as adverse event spontaneous reporting systems and electronic health records) offer an opportunity to mine for new QT-DDIs, but when used individually these datasets have a number of limitations that prevent identification of true signals.
- We present an integrative data science approach that combines mining for latent QT- DDI signals in the FDA Adverse Event Reporting System and retrospective analysis of electrocardiogram lab results in electronic health records at Columbia University Medical Center to identify 26 novel QT-DDIs.

## 1 Introduction

Long QT syndrome (LQTS) is a genetic or acquired change in the electrical activity of the heart that can increase risk of *torsades de pointes* (TdP), a dangerous ventricular tachycardia that can lead to sudden cardiac death [1]. Diagnosed using an electrocardiogram (ECG), LQTS is characterized by a prolonged QT interval and represents an abnormally increased cardiac action potential duration. While the link between QT prolongation and TdP is complex and involves the interplay of multiple factors, a QT interval > 500 ms (versus a normal range of 350 – 440 ms) is nonetheless considered a significant risk for arrhythmogenesis [2].

Since the first reports of TdP in the 1960s [3], mutations in 13 genes coding for cardiac ion channels and their associated proteins have been found to play roles in LQTS [1,4-6].Congenital LQTS can result from mutations that disrupt the *I*_*Ks*_, *I*_*Kr*_, or *I*_*Na*_ ion currents; however, the acquired form of LQTS (which is often drug-induced) is almost exclusively due to block of the HERG channel (*KCNH2*) which plays a role in the *I*_*Kr*_ delayed rectifier potassium current responsible for ventricular repolarization [3]. Drug-induced inhibition of *I*_*Kr*_ was first discovered for the antiarrhythmic quinidine [7], and since then 180 drugs with both cardiac and non-cardiac indications have been found to possess either a known, possible, conditional, or congenital link to dangerously prolonging the QT interval [8]. Terfenadine (an allergy medication) and cisapride (used to treat acid reflux) were withdrawn from the market in 1997 and 2000 respectively for prolonging the QT interval [9], and risk of TdP is now the second leading cause for approved drug withdrawal [2].

Drug-drug interactions (DDIs) such as those between methadone (an analgesic) and quetiapine (an antipsychotic) have also been reported to increase risk for TdP [10]. Despite the increasingly comprehensive resources available to clinicians for linking single drugs to TdP, little remains known about DDIs (QT-DDIs). While the FDA has required clinical studies to assess the effects of drug interactions, it is intractable to prospectively evaluate every possible drug combination. With DDIs thought to play a role in upwards of 9% of adverse events and an increasingly aging population taking multiple drugs concurrently [11,12], there is a pressing need for methods to identify potential interactions.

Molecular mechanism-based approaches such as biological network analysis have been used previously to prioritize drugs with molecular links to LQTS genes, but they remain limited to known drug targets and often only apply to individual drugs [6]. More recent work using machine learning on network data can overcome the requirement for known targets [13]; however, this approach has only been validated for individual drugs.

Observational healthcare datasets such as the FDA Adverse Event Reporting System (FAERS) and electronic health records (EHRs) provide invaluable resources for adverse event (AE) prediction, but their use is tempered by multiple limitations. Spontaneous reporting systems such as FAERS are known to suffer from both reporting bias and sampling variance [14], and methods for mining FAERS traditionally rely on direct evidence between a drug exposure and adverse event (i.e. the number of reports with the drug and AE co-mentioned). While methods have been developed to limit high false positives by correcting for unsubstantiated drug-AE signals [15], this leads to a tradeoff between reducing false positive rates and the ability to actually detect AEs. Direct detection of adverse events falters in the prediction of DDIs, where reporting numbers are often very low. Combined with underreporting of unanticipated or unexpected events with no understood molecular explanation, most DDI signal detection algorithms have had limited success [16,17]. Additionally, AE detection in EHRs can be challenging as such data are often complex, inaccurate, and missing [18]. While use of either dataset alone can thus be problematic for QT-DDI detection, integration of these two sources using data science offers an opportunity for improved performance.

In previous work, we demonstrated that a novel signal detection algorithm could be used for detecting latent signals of previously unknown DDIs for eight severe adverse event classes [19,20]. Importantly, each individual drug in the drug pair had no previously known connection to the AE class of interest. In this study, we introduce an updated pipeline called DIPULSE (Drug Interaction Prediction Using Latent Signals and EHRs) that uses latent signal detection in FAERS to generate an *adverse event fingerprint* for LQTS. This AE fingerprint – trained on individual drugs with a known link to prolonging the QT interval – is then used to predict new QT-DDIs where neither drug alone has a known association to this phenotype. We validate these predictions using ECG lab results in electronic health records (EHRs).

## 2 Methods

A graphical overview of DIPULSE can be found in Figure 1. We implemented the method using Python 2.7.9 and R 3.1.0.

**Figure 1:**
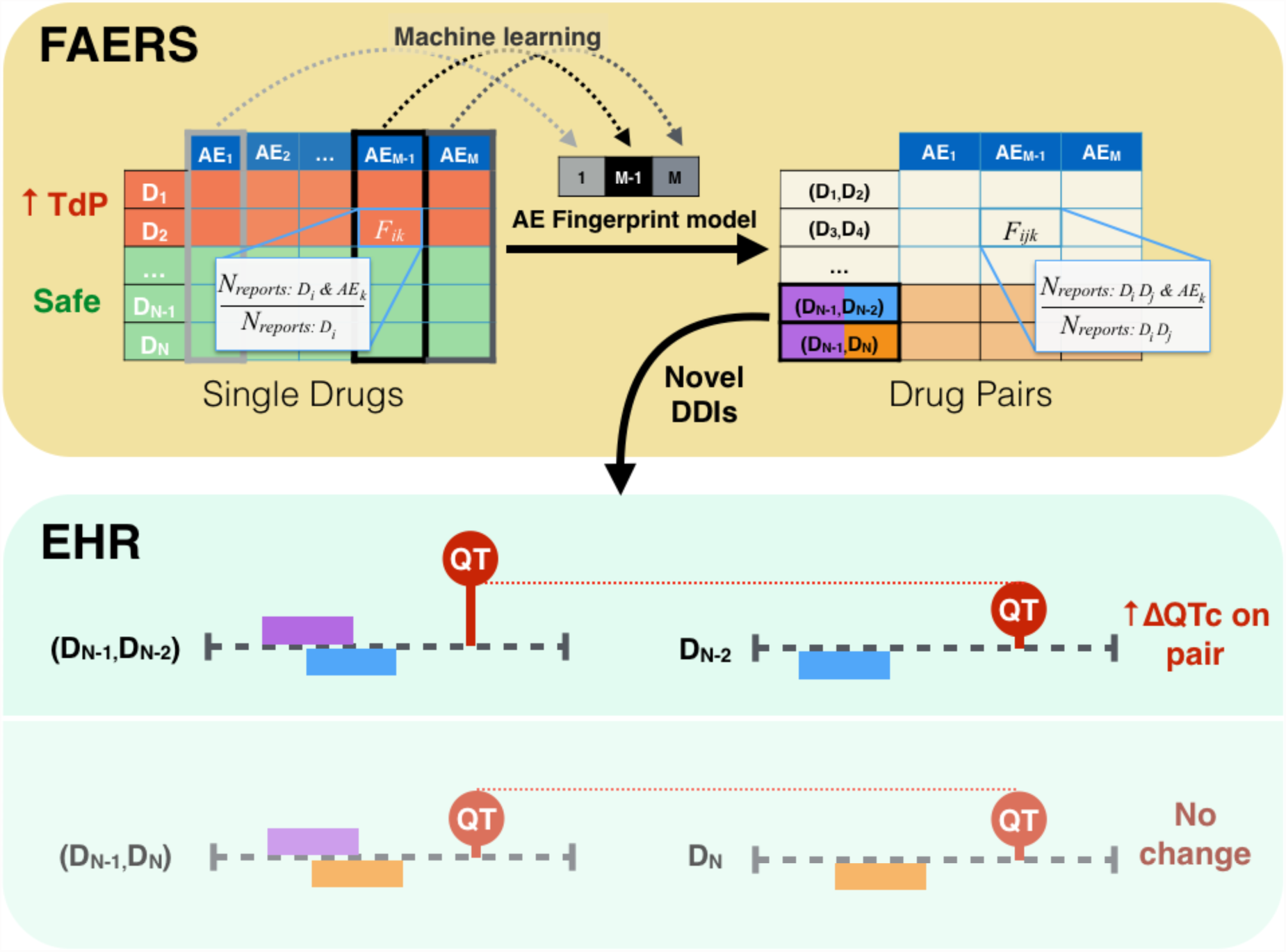
Overview of DIPULSE pipeline. Drug Interaction Prediction Using Latent Signals and EHRs combines mining of the FDA Adverse Event Reporting System (FAERS) and electronic health records (EHRs) to flag novel QT-prolonging drug-drug interactions (DDIs). **FAERS:** We generate an adverse event (AE) reporting frequency table (dimensions *N* drugs by *M* AEs) for single drugs in FAERS. We label a drug as a positive example if it has a known risk of TdP (obtained from CredibleMeds.org). All drugs not found in CredibleMeds are labeled as negative examples. We use machine learning to generate an *AE fingerprint model* that identifies the most predictive subset of features (AE reporting frequencies, *F*_*ik*_) as latent evidence for predicting whether a drug does or does not prolong the QT interval. We then apply this fingerprint model to an independent test data set consisting of a matrix (with AE reporting frequencies *F*_*ijk*_) for drug pairs. We send pairs receiving high classifier probabilities (but where neither individual drug is known to prolong the QT interval) for EHR validation. **EHR:** We validate putative interactions using electrocardiogram lab results in the electronic health records (EHRs) by determining whether patients prescribed a predicted interacting drug pair have increased QTc intervals compared to patients taking either drug alone. In this example patients prescribed drug pair (D_N-1_,D_N-2_) have a significantly increased QT interval compared to patients on either drug alone. This is not observed for drug pair (D_N-1_,D_N_) so it is filtered out. Finally, we perform a confounding analysis to confirm that the significant increase observed in QTc interval is not due to other co-prescribed medications.

### 2.1 Data Sources

We downloaded the intermediate adverse event reporting frequency tables from the OFFSIDES (single drug) and TWOSIDES (drug pair) databases [21]. These tables were generated from 1,851,171 reports in FAERS (corresponding to the first quarter of 2004 to the first quarter of 2009). Each report in FAERS contains the drugs prescribed to the patient, the drug indications, and the observed adverse events. OFFSIDES and TWOSIDES were created by training propensity score matching (PSM) models to match patients exposed to a single drug or drug pair to unexposed controls on the basis of co-prescribed medications and drug indications; an advantage of this approach is that only patients for whom controls could be matched are used for drug safety prediction [21].

An intermediate step in this process is the assembly of adverse event frequency reporting tables for both single drugs and drug pairs as seen in Figure 1, with each row representing a drug and each column one of the AEs in FAERS. For single drugs, the value at a given row and column represents the frequency of reporting *F*_*ik*_, defined as the fraction of reports for drug *i* containing the AE *k*. Similarly, for drug pairs, the reporting frequency *F*_*ijk*_ corresponds to the fraction of reports for drug pair (*i*, *j*) containing the AE *k*. We used the former matrix to train the fingerprint model and the latter for DDI prediction.

As positive controls, we downloaded a list of 180 drugs with known (N=47), possible (N=72), conditional (N=30), or congenital (N=27) risk of TdP from CredibleMeds, an online compendium of drugs associated with LQTS [8]. We also obtained a list of 2,856 critical and significant DDIs from the Veteran Affairs Hospital [22].

To validate our DDI predictions, we used EHR data from Columbia University Medical Center (CUMC). In addition to patient demographics, drugs prescribed, and diagnosis codes, we also used QTc (heart rate-corrected QT interval) values obtained from ECG lab results. The study was approved by the CUMC Institutional Review Board.

### 2.2 Training adverse event fingerprint model

We used the reporting frequencies (*F*_*ik*_) in the frequency table for single drugs as features to train a logistic regression classifier. The binary classifier learns how to weight each feature (an AE reporting frequency, *F*_*ik*_) to best classify a given drug as prolonging the QT interval or not. Training the model requires both positive and negative examples. Our positive examples were all drugs in FAERS with a known risk of TdP (N = 18). As negative controls we selected all drugs in FAERS that do not appear in CredibleMeds (i.e. have no known, possible, conditional, or congenital risk of TdP, N = 562).

Because the number of features (11,305 AEs) is much greater than the number of examples (580 drugs), overfitting of the model to the training data is a concern. To ensure the model generalized to our test data set (drug pairs), we reduced the number of features by using L1 (lasso) regularization. L1 regularization is preferred because it results in sparse models (i.e. most of the feature weights will be driven to zero). We generated five models, each of which contained between 5 - 20 features obtained by varying the regularization strength for the given model. We evaluated these models using 10-fold cross-validation and then re-fit the classifier using only the selected features. The features for each of these models constitute an *adverse event fingerprint* that represents latent evidence for QT interval prolongation.

### 2.3 Predicting novel drug-drug interactions using the fingerprint model

We next applied the QT fingerprint model to an independent test data set consisting of the AE reporting frequencies (*F*_*ijk*_) in the frequency table for drug pairs. The model outputs a probability for a given drug pair to prolong the QT interval. We assessed model performance using two references. In the first, we labeled each drug pair containing a drug known to increase risk of TdP as a positive example; while these may not be bonafide DDIs, they demonstrate the ability of the fingerprint model to “ re-discover” drugs known to prolong the QT interval within the drug pair data. We used this validation to select the optimal fingerprint model. We also performed an additional validation using a list of critical and significant DDIs from the Veteran Affairs Hospital.

As a control, we generated a logistic regression model built using solely direct evidence of QT interval prolongation. There were only five AEs corresponding to QT interval prolongation or TdP, and therefore feature selection was not necessary. We compared the performance of this “ direct-evidence” model against the “ latent” AE fingerprint model using DeLong’s test [23].

To obtain a candidate list of novel DDIs predicted by the fingerprint model, we first removed all drug pairs containing a drug in the CredibleMeds list. We then filtered for all novel predictions found at a classifier probability below a 5% false positive rate (FPR) according to the CredibleMeds evaluation. Finally, to account for the potential of false discoveries we estimated empirical q-values for each drug pair by generating 100 logistic regression models using randomly chosen features. We removed any drug pairs receiving an empirical q-value ≥ 0.01.

### 2.4 Validating novel drug-drug interactions using electronic health records

While the novel DDIs predicted using our signal detection algorithm each contain latent evidence for prolonging the QT interval, ECG values in electronic health records allow us to retrospectively evaluate the effect of these drug pairs (our cases) on QT interval duration compared to either drug alone (our controls). Because QT interval durations differ between males and females [24], we evaluated the effects of a given drug pair on each sex separately.

To obtain cases, we selected patients at New York-Presbyterian Hospital/ Columbia University Medical Center prescribed each drug in a given drug pair within a 7 day period. Patients were also required to have an ECG lab – and corresponding QTc (heart rate-corrected QT interval) – within 36 days of the second drug prescription. For patients with multiple QTc values within this time period, we used the maximum value.

To obtain controls, we selected patients taking whichever individual drug in the pair yielded the greatest median QTc within a 36 day period from drug prescription. We then compared QTc values between cases and controls and assessed significance using a Mann-Whitney U test, correcting for multiple hypothesis testing using Bonferroni’s method.

In order to demonstrate that the predictions being sent for EHR validation were enriched for drug interactions that actually prolonged the QT interval, we ran the above EHR case-control analysis on a set of 10,000 randomly chosen drug pairs found in FAERS. From this set we then randomly sampled with replacement a number of drug pairs equal to that generated by the latent signal detection and counted the number of pairs that had significant increases in QT interval. We repeated this sampling 1,000 times to build an empirical distribution of how many significant results would be expected after EHR analysis by chance alone.

Finally, we performed a confounder adjustment to confirm that the elevated QTc interval on the drug pair was not due to other co-prescribed medications. For each of our sets of cases (patients on a given drug pair) and controls (patients on an individual drug in the pair), we identified possible confounder drugs by counting the number of exposures to each drug prescribed up to 36 days prior. We evaluated each potential confounder by confirming that it was correlated both with the exposure condition and with QTc values. For the former, we determined whether the covariate was more likely to be prescribed with the drug pair compared to the single drug using a Fisher’s exact test; for the latter we compared the QTc values for patients exposed to the covariate versus those unexposed using a Mann-Whitney U test. Both of these evaluations were performed using a Bonferroni correction for multiple hypothesis testing. We collected all drug covariates that passed these two requirements and assessed their significance (for males and females separately) using an analysis of covariance (ANCOVA). To obtain the final list of validated novel DDIs, we only kept those results (drug pairs for a given sex) receiving significant ANCOVA p-values (P < 0.05) for the DDI.

## 3 Results

### 3.1 QT fingerprint model significantly outperforms model built using only direct evidence

Of the five fingerprint models evaluated, we found that the model containing 13 features achieved the best performance for drug pair data (AUC = 0.69 using pairs containing a known CredibleMeds drug) (Supplementary Figure 1); see Table 1 for the list of features that constitute the QT adverse event fingerprint. Importantly, the QT fingerprint model significantly outperformed the model built using direct evidence as evaluated by both the CredibleMeds (P = 9.982E-10) and Veteran Affairs (P = 9.501E-09) drug pair standards (Figure 2). After filtering using both empirical q-values and the 5% FPR cutoff we obtained 1,310 putative novel DDIs to be validated in the EHR.

**Table 1:** Features in QT fingerprint model.

| Adverse Event | Weight |
| --- | --- |
| atrial fibrillation | 0.5 |
| drug interaction | 0.37 |
| bradycardia | 0.3 |
| asystole | 0.27 |
| tachycardia ventricular | 0.27 |
| electrocardiogram QT prolonged | 0.25 |
| completed suicide | 0.23 |
| torsade de pointes | 0.22 |
| agitated | 0.06 |
| heart attack | -0.22 |
| accident | -0.23 |
| abdominal discomfort | -0.3 |
| drug ineffective | -0.3 |

### 3.2 EHR validation and confounder analysis confirms novel drug interactions prolonging the QT interval

Our EHR evaluation yielded 91 results (drug pairs for males and/ or females) that had significantly increased QTc intervals on the drug pair compared to either drug alone (Supplementary Figure 2). This number of results was significantly greater than for randomly generated input to the EHR validation (P < 0.001) (Supplementary Figure 3). After confounder analysis we obtained 32 results (corresponding with 26 distinct drug pairs) that represent validated novel DDIs that increase risk of acquired LQTS (Table 2).

**Table 2:** List of novel DDIs generated by DIPULSE and validated in the EHR.

| Drug 1 | Drug 2 | Control | Sex | Estimate | P | Mean QTc cases | Mean QTc controls | ΔQTc (ms) | # Cases | # Controls |
| --- | --- | --- | --- | --- | --- | --- | --- | --- | --- | --- |
| octreotide | lactulose | *octreotide* | M | 74.9 | 0.0008 | 486 | 455 | **31** | 324 | 535 |
| aspirin | rifampicin | *rifampicin* | M | 57.6 | 6.9E-07 | 469 | 442 | **27** | 177 | 373 |
| mupirocin | nystatin | *mupirocin* | M | 37.4 | 0.00041 | 479 | 452 | **27** | 394 | 1713 |
| aspirin | bacitracin | *bacitracin* | M | 20.9 | 0.02309 | 461 | 439 | **22** | 379 | 814 |
| metoprolol | fosphenytoin | *metoprolol* | M | 54.3 | 1.4E-11 | 462 | 441 | **21** | 429 | 20672 |
| gentamicin | ipratropium bromide | *ipratropium bromide* | M | 38.7 | 0.00053 | 474 | 454 | **20** | 370 | 9251 |
| galactose | vancomycin | *galactose* | M | 26.5 | 2.00E-16 | 465 | 445 | **20** | 5486 | 12121 |
| metolazone | vancomycin | *metolazone* | M | 36.2 | 0.02691 | 492 | 475 | **17** | 337 | 654 |
| N-acetylcysteine | vancomycin | *vancomycin* | M | 20.4 | 6E-06 | 467 | 451 | **16** | 2152 | 8209 |
| heparin | diltiazem | *heparin* | M | 10.2 | 0.0323 | 457 | 442 | **15** | 1781 | 28533 |
| morphine | mupirocin | *mupirocin* | M | 25.7 | 0.0006 | 464 | 450 | **14** | 1183 | 920 |
| N-acetylcysteine | lorazepam | *N-acetylcysteine* | M | 24.4 | 2.8E-07 | 463 | 449 | **14** | 1326 | 5573 |
| heparin | digoxin | *digoxin* | M | 21.6 | 2.9E-07 | 460 | 446 | **14** | 1986 | 1158 |
| ceftriaxone | simvastatin | *ceftriaxone* | M | 26.7 | 0.01716 | 457 | 444 | **13** | 489 | 4768 |
| cefazolin | meperidine | *cefazolin* | M | 17.3 | 3.5E-11 | 450 | 437 | **13** | 1858 | 8362 |
| metoprolol | vancomycin | *vancomycin* | M | 13.3 | 0.00391 | 461 | 449 | **12** | 4108 | 6257 |
| N-acetylcysteine | nystatin | *nystatin* | M | 23.7 | 0.00588 | 460 | 449 | **11** | 890 | 3439 |
| atorvastatin | vitamin K1 | *vitamin K1* | M | 14.5 | 0.03336 | 472 | 462 | **10** | 516 | 3654 |
| ceftriaxone | lansoprazole | *ceftriaxone* | M | 23.5 | 0.00103 | 454 | 444 | **10** | 702 | 4554 |
| N-acetylcysteine | ipratropium bromide | *ipratropium bromide* | M | 13.8 | 0.00026 | 462 | 452 | **10** | 2467 | 7093 |
| lidocaine | meperidine | *lidocaine* | M | 23.2 | 0.00258 | 457 | 448 | **9** | 319 | 4785 |
| aspirin | mupirocin | *mupirocin* | M | 26.3 | 1.3E-08 | 460 | 451 | **9** | 1448 | 666 |
| galactose | aspirin | *galactose* | M | 13.7 | 5.6E-11 | 453 | 447 | **6** | 9879 | 7802 |
| cortisol | vitamin K1 | *vitamin K1* | F | 84.8 | 0.00012 | 476 | 456 | **20** | 556 | 2767 |
| mupirocin | vancomycin | *vancomycin* | F | 77.8 | 3.2E-06 | 473 | 454 | **19** | 566 | 7450 |
| aspirin | mupirocin | *mupirocin* | F | 28.0 | 0.01017 | 470 | 452 | **18** | 686 | 477 |
| galactose | vancomycin | *galactose* | F | 29.4 | 6E-06 | 466 | 448 | **18** | 4138 | 10724 |
| hydromorphone | pravastatin | *pravastatin* | F | 13.8 | 9.1E-07 | 462.5 | 447 | **15.5** | 140 | 908 |
| cefazolin | meperidine | *cefazolin* | F | 31.8 | 3.6E-06 | 453 | 438 | **15** | 818 | 6738 |
| metoprolol | vancomycin | *vancomycin* | F | 15.4 | 1.5E-06 | 464 | 450 | **14** | 2879 | 5111 |
| heparin | digoxin | *heparin* | F | 23.9 | 0.00062 | 455 | 445 | **10** | 1551 | 27277 |
| N-acetylcysteine | lorazepam | *N-acetylcysteine* | F | 32.1 | 0.00042 | 460 | 453 | **7** | 1052 | 3681 |

**Figure 2:**
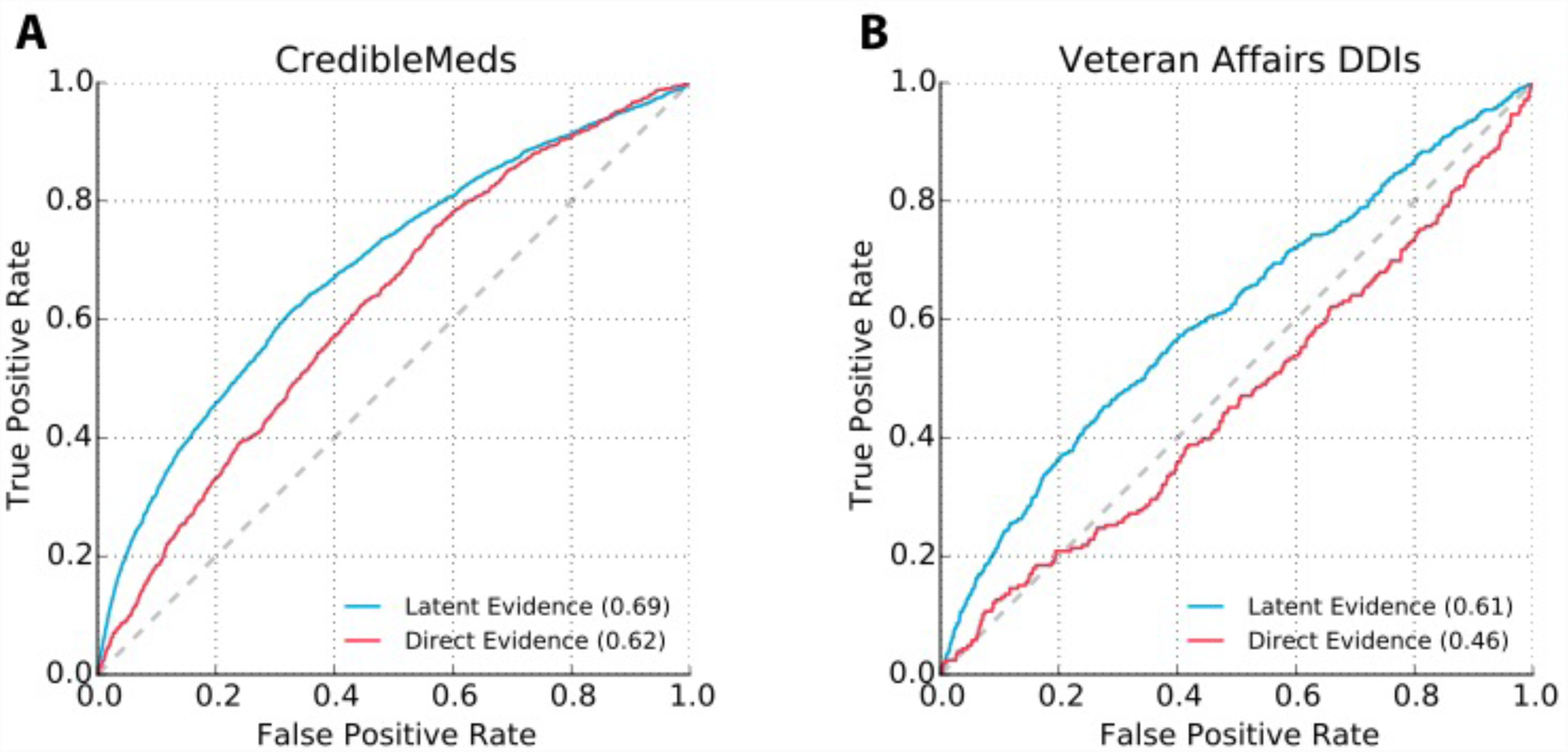
Receiver operating characteristic curves for adverse event fingerprint model and direct evidence control. **(A)** Model validation was performed by labeling drug pairs containing a drug with known increased risk of TdP as positive examples. We compared the performance of a model built using latent evidence (AE fingerprint model) to a control model using only direct evidence of QT prolongation. **(B)** A second evaluation performed using a list of critical and significant DDIs from the Veteran Affairs Hospital in Arizona. For both validations, the AE fingerprint model significantly outperforms the model built solely with direct evidence.

The greatest increase in mean QTc (31 ms) was for octreotide (a somatostatin analog used to lower growth hormone levels) and lactulose (given to treat constipation) compared to octreotide alone (P = 8.02E-4) in males. For females, co-prescription of cortisol and vitamin K1 was associated with a 20ms increase in mean QTc compared to vitamin K1 alone (P = 1.23E- 4). A complete list of retrospectively validated interactions and the number of patients in the case and control groups can be found in Table 2.

## 4 Discussion

Drug-induced LQTS and its potential for fatal arrhythmia (TdP) make this disorder of critical importance both to drug discovery and pharmacovigilance; indeed, an important step in the drug development process is confirming that the lead compound does not significantly block the HERG channel which contributes to TdP [2]. However, the inability to prospectively identify this risk is highlighted by the increasing number of drugs found to increase risk for TdP [8]. Even more difficult to detect are drug-drug interactions that contribute to LQTS, as experimental evaluation of all possible QT-DDIs is not feasible and traditional methods for mining observational data are poorly equipped to handle low reporting numbers and high false positive rates. While analyses of spontaneous reporting systems (such as FAERS) and EHRs alone have many limitations, integrative approaches to incorporate multiple dimensions of observational data can allow for identification of true QT-DDI signals. We demonstrate the applicability of this data science approach in this study by identifying latent signals of LQTS in FAERS and validating these novel QT-DDI predictions, retrospectively, using EHRs.

While most drugs prolong the QT interval by interacting with the HERG channel, the clinical data used in this analysis do not permit a mechanistic explanation for the synergistic effects of the identified DDIs. Electrophysiology experiments to directly assay the effect of individual drugs and drug pairs on HERG channel activity can provide further evidence for and molecular mechanisms of these effects [2]. Importantly, QTc correction formulas still used today were developed in 1920 and are known to be inaccurate when heart rate changes occur outside the baseline range used to define the formula [2]. As such, drugs that do not directly affect ventricular repolarization but instead alter the patient’s heart rate may be incorrectly attributed to increasing the QTc. It is possible that some of the interactions we identified were confounded by this complexity. This limitation highlights the need for experimental validation of our QT-DDI predictions to directly assess HERG channel block or effects on other ion channels.

While cases of drug-induced LQTS have predominantly been found to be due to blocking of *I*_*Kr*_, we do not discount the possibility for other potential mechanisms of these QT- DDIs. Biological network analysis [6,13] may be useful to identify other proteins, in addition to or instead of HERG, that are affected by these drugs.

## 5 Conclusion

In this study we have developed and validated DIPULSE, an automated integrated pipeline for flagging novel drug-drug interactions that can prolong the QT interval using data both from spontaneous reporting systems (FAERS) and electronic health records. By identifying latent signals of QT interval prolongation, the method is able to overcome some of the limitations in mining for drug-drug interactions. The method significantly outperforms DDI detection using solely direct evidence for QT prolongation. This study highlights the utility of integrative data science approaches in mining for new and potentially fatal DDIs.

## Acknowledgements

T.L. is supported by a training grant from the National Institute of General Medical Sciences (T32GM082797). T.L. and N.P.T. are supported by the National Institute of General Medical Sciences (R01GM107145). K.J.S. and R.S.K. are supported by the National Institute of Health (5R01GM109762-02). T.L., K.J.S., R.S.K, and N.P.T. declare that they have no conflict of interest. R.L.W. is an uncompensated officer of the nonprofit organization AZCERT.org, which is supported by FDA HHSF223201400189C and which maintains the the website www.CredibleMeds.org utilized in this study; he declares no conflict of interest We would like to thank Sam Roe for thoughtful discussions about the manuscript.

## Author Contributions

T.L. and N.P.T. designed the research, performed the research, and analyzed the data. K.J.S., R.S.K., and R.L.W. contributed new reagents/analytical tools. T.L. and N.P.T. wrote the article.

## Supplementary Figures

**Supplementary Figure 1:**
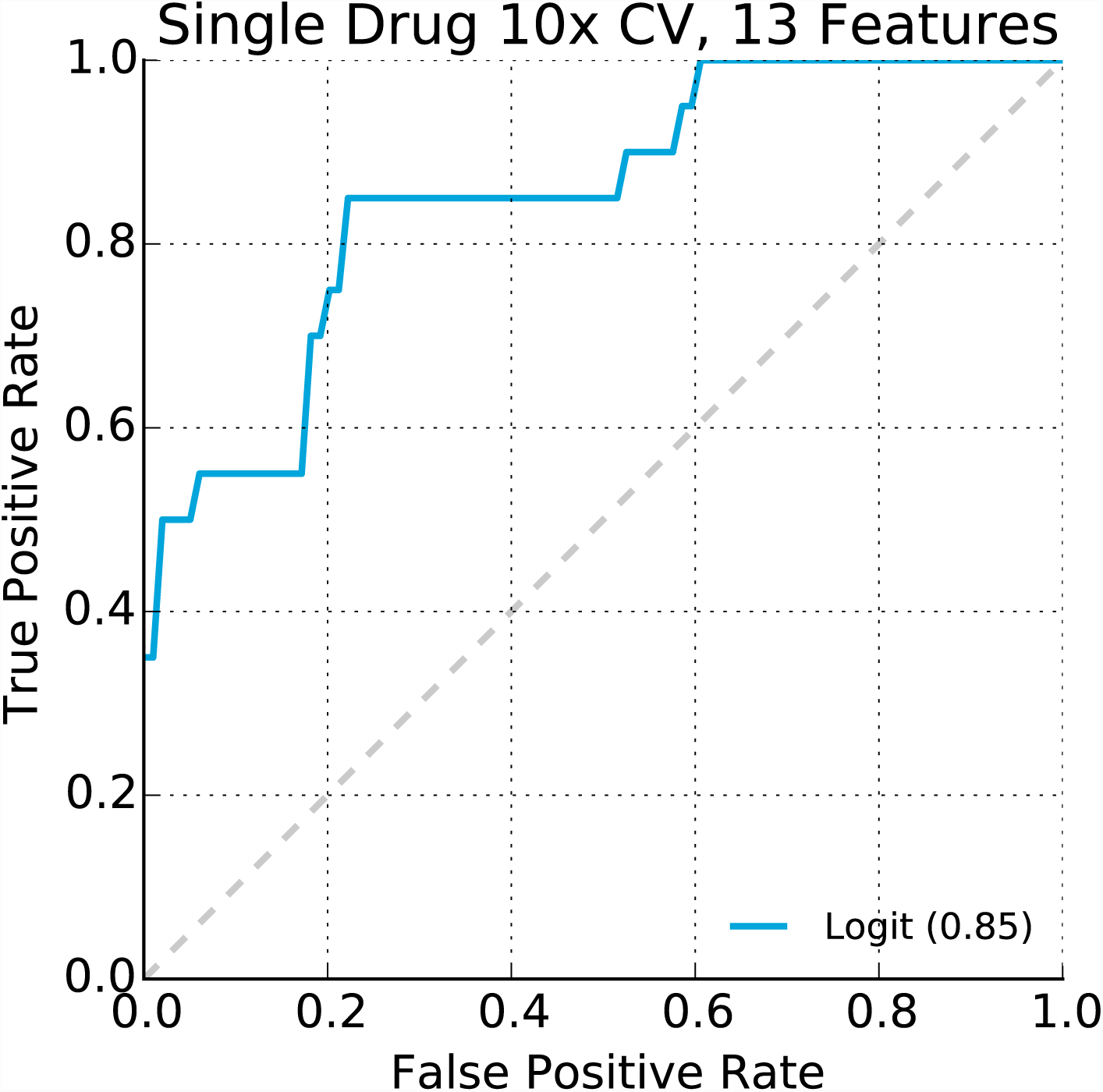
Receiver operating characteristic curve for QT fingerprint model applied to single drugs. As part of building models based on latent evidence, we confirmed that the model could correctly classify the single drugs known to increase risk of TdP in our training set using 10-fold cross-validation. The model generated with 13 features achieved an area under the ROC curve (AUROC) of 0.85.

**Supplementary Figure 2:**
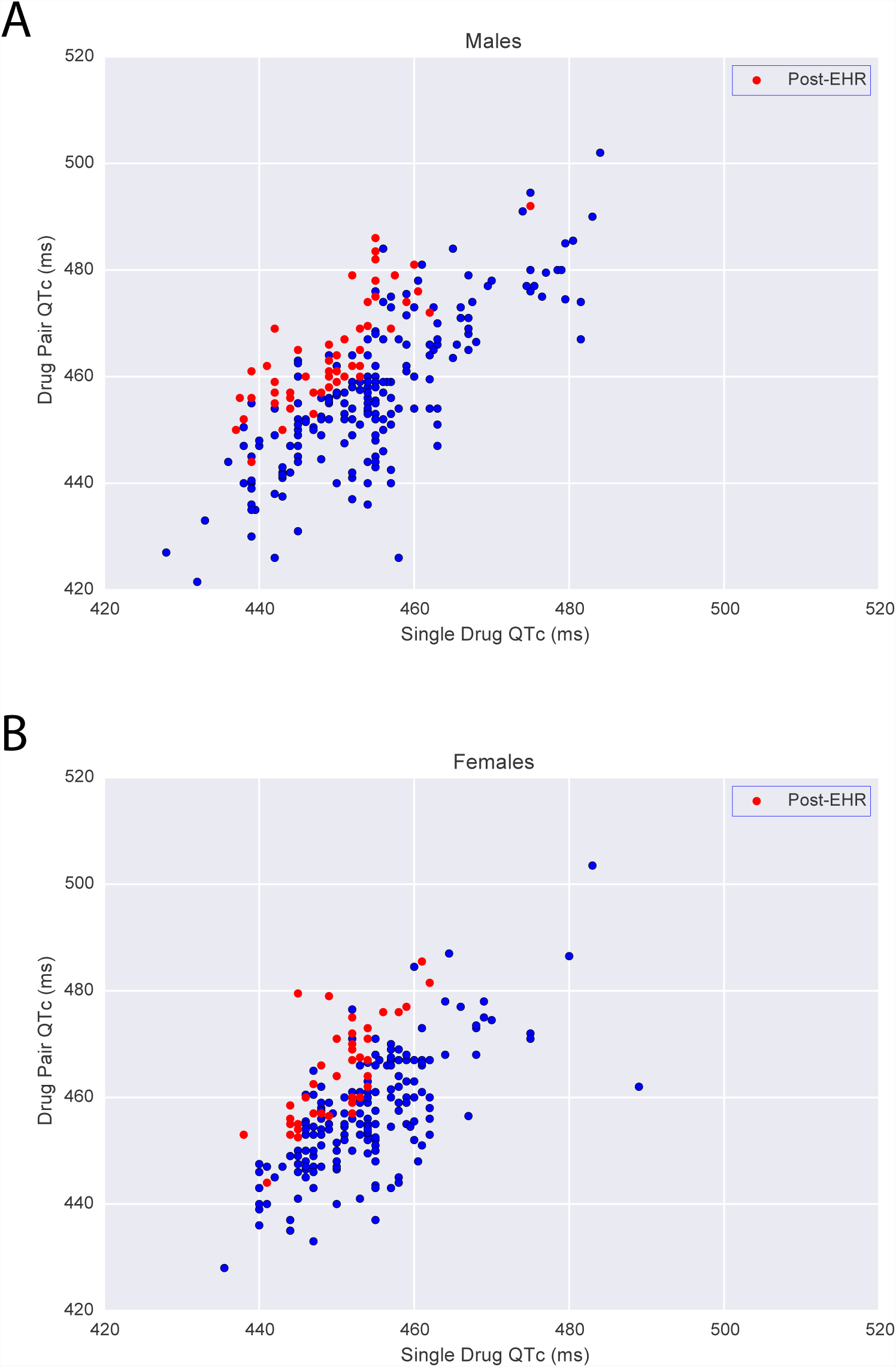
Scatter plot comparing mean QTc intervals (in milliseconds) on single drug (x-axis) and combination therapy (y-axis). **(A)** Results for males. **(B)** Results for females. In both panels a minimum of 50 patients on the drug pair was necessary for inclusion in the plot. A red circle indicates a drug pair that had significantly increased QTc compared to the single drug control in the EHR analysis.

**Supplementary Figure 3:**
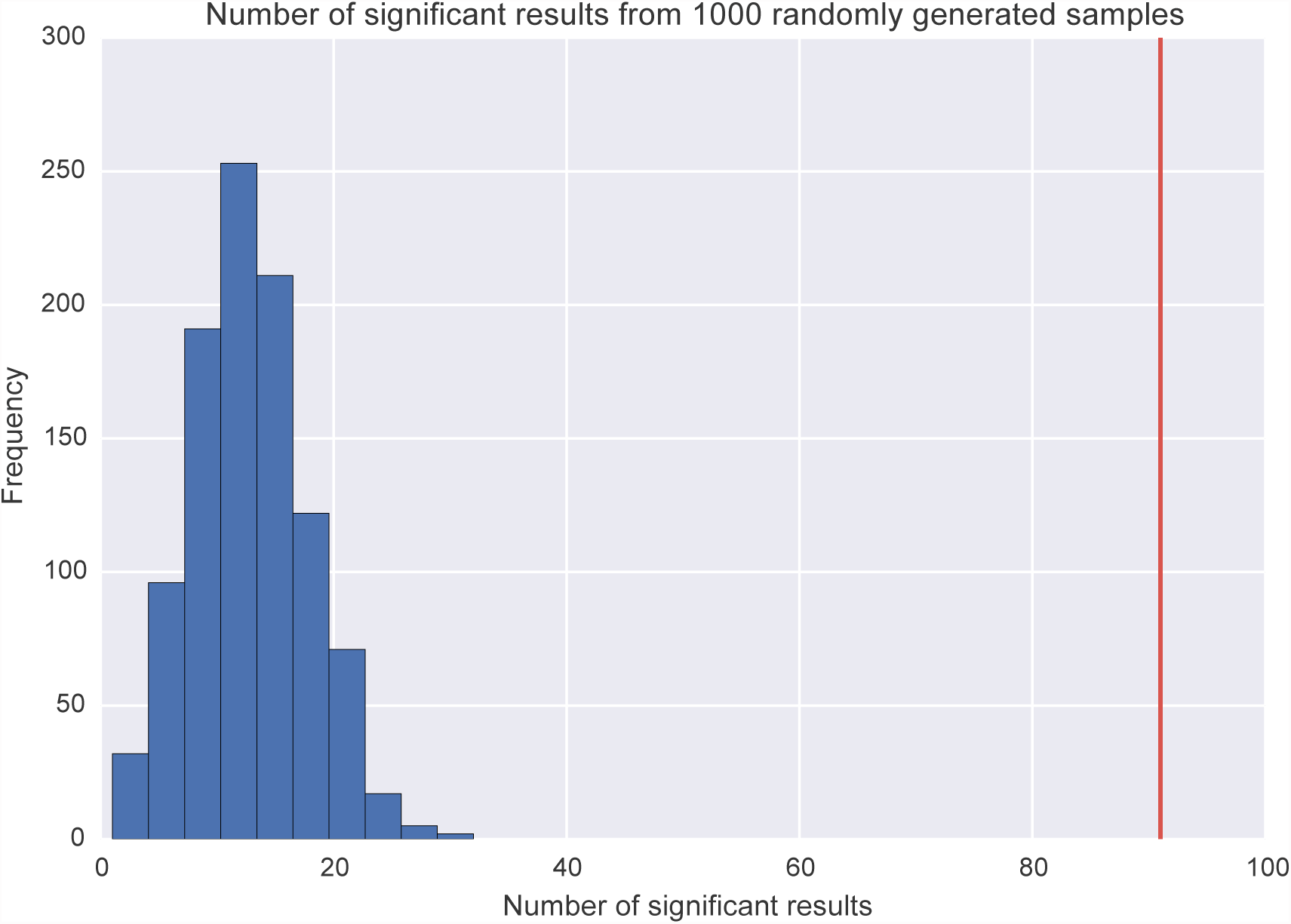
DIPULSE generates significantly more true predictions than would be expected by chance alone. DIPULSE generated 1,310 putative DDIs for EHR analysis, of which 91 results (drug pair and a given sex) were found to be significant (red line). After running 10,000 random drug pairs from FAERS through the EHR case-control analysis, we sampled with replacement 1,000 randomly generated samples of 1,310 pairs and counted the number of significant results to build an empirical distribution (blue). DIPULSE significantly enriched for drug interactions that actually prolong the QT interval (P < 0.001).

## References

1. Roden DM. Clinical practice. Long-QT syndrome. N Engl J Med. 2008;358:169–76.

2. Fermini B, Fossa AA. The impact of drug-induced QT interval prolongation on drug discovery and development. Nat Rev Drug Discov. 2003;2:439–47.

3. Kannankeril P, Roden DM, Darbar D. Drug-Induced Long QT Syndrome. Pharmacological Reviews. 2010;62:760–81.

4. Marx SO, Kurokawa J, Reiken S, Motoike H., D’Armiento J, Marks AR, et al. Requirement of a macromolecular signaling complex for β adrenergic receptor modulation of the KCNQ1-KCNE1 potassium channel. Science. 2002.

5. Moss AJ, Kass RS. Long QT syndrome: from channels to cardiac arrhythmias. J. Clin. Invest. 2005;115:2018–24.

6. Berger SI, Ma’ayan A, Iyengar R. Systems Pharmacology of Arrhythmias. Science Signaling. 2010;3:ra30–0.

7. Roden DM, Woosley RL, Primm RK. Incidence and clinical features of the quinidine-associated long QT syndrome: implications for patient care. Am. Heart J. 1986;111:1088–93.

8. Woosley RL, Romero K. Assessing cardiovascular drug safety for clinical decision-making. Nat Rev Cardiol; 2013;10:330–7.

9. Woosley RL, Chen Y, Freiman JP, Gillis RA. Mechanism of the cardiotoxic actions of terfenadine. JAMA. 1993.

10. Uehlinger C, Crettol SV, Chassot P, Brocard M, Koeb L, Brawand-Amey M, et al. Increased (R)-Methadone Plasma Concentrations by Quetiapine in Cytochrome P450s and ABCB1 Genotyped Patients. J Clin Psychopharmacol. 2007;27:273–8.

11. Leape LL, Bates DW, Cullen DJ, Cooper J. Systems analysis of adverse drug events. JAMA. 1995.

12. Hajjar ER, Cafiero AC, Hanlon JT. Polypharmacy in elderly patients. Am J Geriatr Pharmacother. 2007;5:345–51.

13. Lorberbaum T, Nasir M, Keiser MJ, Vilar S, Hripcsak G, Tatonetti NP. Systems pharmacology augments drug safety surveillance. Clin Pharmacol Ther. 2015;97:151–8.

14. Bate A, Evans SJW. Quantitative signal detection using spontaneous ADR reporting. Pharmacoepidem. Drug Safe. 2009;18:427–36.

15. Szarfman A, Machado SG, O’Neill RT. Use of screening algorithms and computer systems to efficiently signal higher-than-expected combinations of drugs and events in the US FDA’s spontaneous reports database. Drug Saf. Springer; 2002;25:381–92.

16. Norén GN, Sundberg R, Bate A, Edwards IR. A statistical methodology for drug–drug interaction surveillance. Statist. Med. 2008;27:3057–70.

17. Harpaz R, Chase HS, Friedman C. Mining multi-item drug adverse effect associations in spontaneous reporting systems. BMC Bioinformatics. BioMed Central Ltd; 2010;11:S7.

18. Hripcsak G, Albers DJ. Next-generation phenotyping of electronic health records. Journal of the American Medical Informatics Association. 2013;20:117–21.

19. Tatonetti NP, Denny JC, Murphy SN, Fernald GH, Krishnan G, Castro V, et al. Detecting drug interactions from adverse-event reports: interaction between paroxetine and pravastatin increases blood glucose levels. Clin Pharmacol Ther. 2011;90:133–42.

20. Tatonetti NP, Fernald GH, Altman RB. A novel signal detection algorithm for identifying hidden drug-drug interactions in adverse event reports. Journal of the American Medical Informatics Association. 2012;19:79–85.

21. Tatonetti NP, Ye PP, Daneshjou R, Altman RB. Data-Driven Prediction of Drug Effects and Interactions. Science Translational Medicine. 2012;4:125ra31–1.

22. Olvey EL, Clauschee S, Malone DC. Comparison of critical drug-drug interaction listings: the Department of Veterans Affairs medical system and standard reference compendia. Clin Pharmacol Ther. 2010;87:48–51.

23. Robin X, Turck N, Hainard A, Tiberti N, Lisacek F, Sanchez J-C, et al. pROC: an open-source package for R and S+ to analyze and compare ROC curves. BMC Bioinformatics. 2011;12:77.

24. Rautaharju PM, Zhou SH, Wong S, Calhoun HP, Berenson GS, Prineas R, et al. Sex differences in the evolution of the electrocardiographic QT interval with age. The Canadian journal of cardiology. 1992;8:690–5.

